# Stochastic methods for inferring states of cell migration

**DOI:** 10.1101/249656

**Authors:** R.J. Allen, C. Welch, N. Pankow, K. Hahn, Timothy C. Elston

**Author notes:** Current Address: Internal Medicine Research Unit, Pfizer Inc Cambridge, MA 02139. Current Address: Department of Otolaryngology/Head and Neck Surgery University of North Carolina School of Medicine, Chapel Hill NC 27599. To whom Correspondences should be addressed: Department of Pharmacology University of North Carolina at Chapel Hill Genetic Medicine Building, Campus Box 7260 Chapel Hill, NC 27599-7260.

## Abstract

Cell migration refers to the ability of cells to translocate across a substrate or through a matrix. To achieve net movement requires spatiotemporal regulation of the actin cytoskeleton. Computational approaches are neceary to identify and quantify the regulatory mechanisms that generate directed cell movement. To address this need, we developed computational tools, based on stochastic modeling, to analyze time series data for the position of randomly migrating cells. Our approach allows parameters that characterize cell movement to be efficiently estimated from time series data. We applied our methods to analyze the random migration of Mouse Embryonic Fibroblasts (MEFS). Our analysis revealed that these cells exist in two distinct states of migration characterized by differences in cell speed and persistence. Further analysis revealed that the Rho-family GTPase RhoG plays a role in establishing these two states. An important feature of our computational approach is that it provides a method for predicting the current migration state of an individual cell from time series data. Using this feature, we demonstrate that HeLa cells also exhibit two states of migration, and that these states correlate with differences in the spatial distribution of active Rac1.

## Introduction

The ability of cells to move is essential to many biological processes, such as tissue development, the immune response and wound healing [1]–[3]. Anomalous cell migration plays a role in diseases, such as cancer and atherosclerosis [2], [4]–[6]. During cell migration, intracellular signaling networks tightly control the spatiotemporal dynamics of the cytoskeleton. In particular, the Rho family of small GTPases has been implicated in membrane protrusion, adhesion, contraction and de-adhesion, all steps necessary for cell migration [7]–[12].

During random cell migration, in which cells do not experience directional environmental cues, cells move in a persistent manner, but with significant variability in their direction and speed. Therefore, methods for quantifying cell movement that take into account the stochastic nature of this phenomenon are needed. Previous studies have analyzed cell migration in terms of quantitative metrics such as the mean squared deviation in cell position, which can be linked to both speed and persistence [13]–[16]. Additionally, it has been suggested that fractional diffusion models are required to accurately describe cell movement [13]. We used stochastic modeling to develop tools for quantifying cell migration such that it can be characterized in terms of biologically relevant parameters. In our approach, the motion of cells is assumed to follow a 2D random walk with persistence. A related method that takes into account the probability of turning and contains a parameter related to persistence also has been applied to analyze random cell migration [17]. An important distinction of our approach is that our model allows for the possibility of multiple states of migration, distinguished by differences in speed and persistence. This feature allowed us to determine that Mouse Embryonic Fibroblasts (MEFS) exist in two distinct states during random migration. Knock down of the Rho-GTPase RhoG, suggests that this protein plays an important role in establishing the two states. We next demonstrated how our method allows the migration state of cell to be predicted from time series data. HeLa cells expressing a Rac1 biosensor revealed that these cells also undergo two states of migration, and that these two states correlate with the number of active Rac1 foci.

## Results

### Preliminary Analysis

To develop our methods, we collected data sets that consisted of time series for the x and y coordinates of the cell centroids of randomly migrating MEF cells (Fig. 1A and B). We chose MEFs because these cells show persistent migration in the absence of directional cues. As an initial analysis of the data, we computed the average persistence of cell movement defined as 
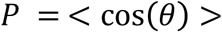
, where *θ* is the change in the direction of cell movement between measurements (Fig. 1C) and the angular bracket denotes averaging over cell tracks. If *θ* is uniformly distributed, then the motion of the cell lacks persistence and P = 0. This behavior would be consistent with a pure random walk (diffusive motion). For values of P greater than zero, the movement of the cell shows persistence, with a value of 1 indicating motion in a straight line. Combining the cell tracks for individual cells, produced a value of P = 0.43. This value is consistent with cells that show highly persistent motion. We also generated a histogram from all the Δx and Δy displacements and empirically calculated cumulative density functions for each (**Sup Fig. 1**). For purely diffusive motion, the step sizes follow a Gaussian distribution. However, the experimentally measured distribution was found to deviate significantly from Gaussian (**Sup Fig. 1**).

**Figure 1.**
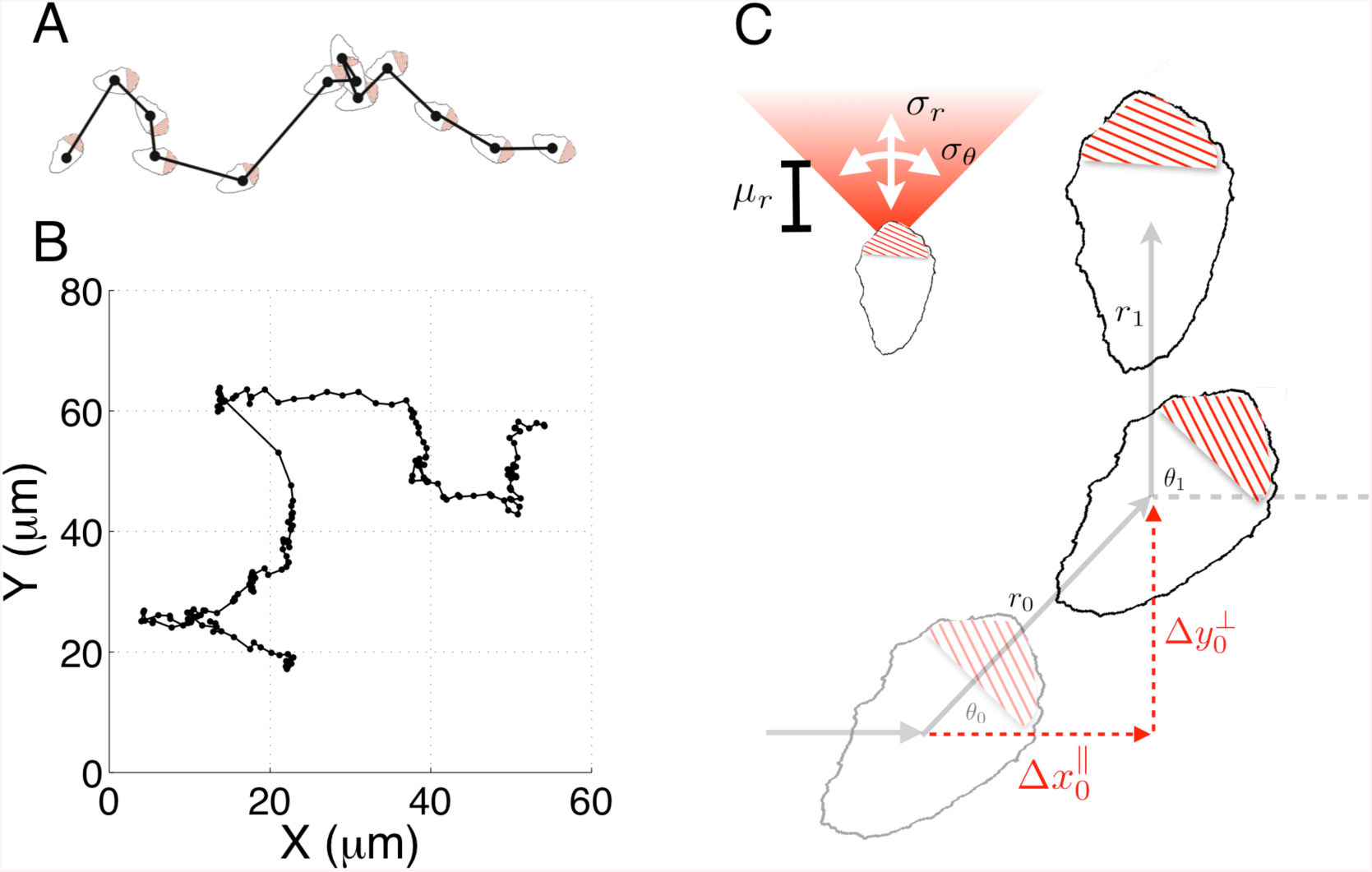
**A** Tracking and modeling of random cell migration. Cell tracks are the result of recording the geometric center of the cell over a time-course. **B** Example track resulting from tracking the cell centroid at 5 minute time intervals (black dots). **C** A simple model of migration, where over a time step the cell moves a distance *μ_r_* on average (with variance 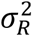), on average the cell continues to move in the same direction (with variance 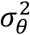). We apply this model by using an analytical expression for the probability density function of the distance travelled in the direction of the prior orientation, 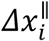.

### A stochastic model for cell migration

Our preliminary cell track analysis led us to model cell movement as a 2D random walk with persistence (Fig. 1C). In our model, for each time interval i, the distance, *r_i_*, traveled by a cell and the angle, *θ*_*i*_, through which the cell moves are considered random variables. The random variable *r_i_* is taken to have a Gaussian distribution characterized by mean *μ_R_*, and variance 
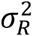
. We allowed for negative values of *r_i_* to account for the scenario in which a cell maintains its direction of polarization, but its centroid moves in a rearward direction. The directional angle *θ*_*i*_, is also taken to have a Gaussian distribution with variance 
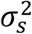
, and centered on the value of the previous angle *θ*_*i*-1_. Small values of 
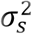
 correspond to highly persistent migration. For large values of 
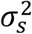
 the new direction becomes uniformly distributed on the interval [-π, π] and the model represents a purely diffusive process.

It is not possible to tell from cell track data if changes in *θi* of magnitude greater than π/2 resulted from large deviations in orientation or negative *r_i_*. Thus, the probability distribution for these variables cannot be constructed unambiguously from the cell track data. To overcome this difficulty, we performed a change of variables from (*r_i_*, *θ*_*i*_) to 
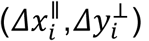
, where these new variables correspond to changes in the centroid’s position during the i^th^ time interval that are parallel and perpendicular to the direction of the previous step (Fig. 1C). An important feature of the model is that analytical expressions for the probability density functions (pdf) of 
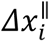
 and 
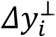
 can be found (Supplemental Material) allowing estimation of model parameters from experimental data to be performed in a computationally efficient manner. These co-ordinates explicitly handle the degeneracy described above, because in these co-ordinates all possibilities that could have led to a given observation are considered. If cells show persistent motion, 
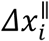
 has a positive mean value. Also, if there are no external cues in the experiments to define a preferred direction of motion, 
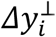
 is symmetric about zero. Therefore, the distribution for 
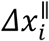
 is more informative, and we use it to compare the experimental results with the model’s behavior. It is possible to simultaneously fit the 
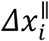
 and 
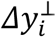
 distributions, but this comes at an increased computational cost. As a consistency check, after performing parameter estimation, we verify that the model accurately captures the 
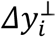
 distribution. If the model failed this consistency test, we could repeat the parameter estimation using both distributions. However, this was not required for any of the cases considered here.

We used a Monte Carlo method based on the Metropolis algorithm to perform parameter estimation. This was followed by local optimization algorithms to identify parameters associated with the global minimum error between the model and data (Supplementary Material). To test the accuracy and efficiency of this method, we benchmarked our approach using data generated from computational simulations of the stochastic model (**Suppl.** Fig. 2). Having validated our computational methods, we next attempted to fit the model to the experimentally measured distribution. However, the model did not generate a good fit to the experimental data (**Suppl.** Fig. 3). In particular, we found that the model could not capture a second mode observed in the distribution for 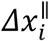.

### A multistate model for cell migration

Further inspection of the cell tracks, and the distribution for 
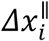
, suggested that individual cells might exist in different modes of migration, distinguished by differences in the speed and persistence. Therefore, we expanded our model to allow for different states of migration. That is, we hypothesized that at any given time a migrating cell is in one of n states denoted by S_i_, with *i* ∈ {1 … *n*}. Each state is characterized by the parameters 
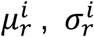
 and 
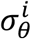
. The additional parameters, *α^i^*, denoting the fraction of time spent in state *i*, are required to fully specify the model. Since 
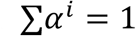
, in the two-state case the total number of parameters is 7. Note that if a two-state model is fit to data consisting of only a single state, then we expect our Monte Carlo method to produce parameter sets in which *α*^1^ takes on values of 0 or 1, or 
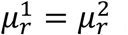
,

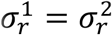
 and 
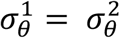
. The extended model is essentially a mixture model, which is itself a reduced hidden Markov model under the assumption that the probabilities of transitioning between states are independent and identically distributed. We again used simulated data to validate the accuracy and efficiency of our Monte Carlo method when multiple states are considered (**Suppl.** Fig. 4).

Using the multi-state model, the Monte Carlo method produced a good fit to the 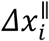 distribution (Fig. 2A). To assess the accuracy of our parameter estimates we used confidence-interval profiling [18]. To determine acceptable values for the sum of the squared errors (SSE) we boot-strapped the original datasets to assess plausible differences in our observed distributions should we repeat the experiments (Supplementary Material). The results of this analysis provides a measure of the confidence that should be placed on each estimated parameter value (**Suppl. Fig. 5**). Of particular interest is the parameter α which represents the fraction of time in each state. For this parameter, our analysis provided further evidence that the data were not consistent with a single state model (α = 1 or 0). The best fits were achieved with α = 0.12. We confirmed that the model also captured the distributions for 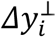 (**Suppl. Fig. 6**). The results of our analysis suggest that randomly migrating MEFs exist in one of two states. About 12% of time these cells are in a state with a well-defined characteristic step of ~3 μm (State 1 - blue distribution in Fig. 2A left inset) and an angular distribution with 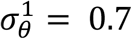. In the second state, the step size is highly variable (State 2 – red distribution Fig. 2A right inset) and the motion is less persistent 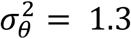. For completeness, we also show the distribution for the angle θ (Fig. 2B).

**Figure 2.**
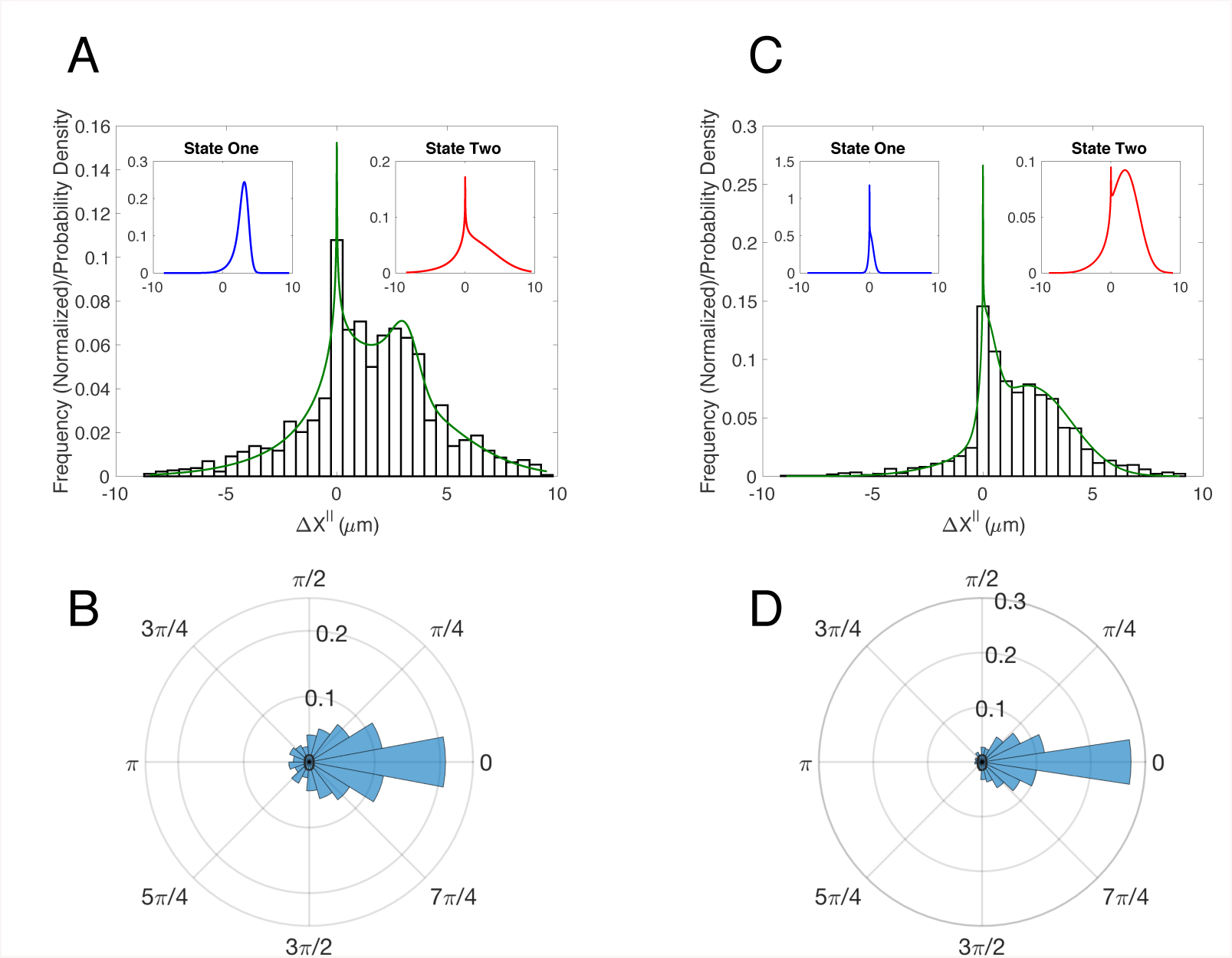
Fitting the model to MEF WT cells and MEF RhoG KD cells identified two states in each case. **A** MEF WT **B** MEF RhoG KD. In **A** and **B**: Observed data (open bars), model fit (green) and individual pdfs for state one and state two (insets). **C** MEF WT. Observed angular distribution for angle turned at each time step, *Δθ* **D** MEF RhoG KD. Observed angular distribution for angle turned at each time step, *Δθ*.

### RhoG’s role in migration

It has long been appreciated that the canonical Rho-GTPases RhoA, Rac1, and Cdc42 play important roles in cell migration. However, the role of RhoG in migration is less well studied. To determine if RhoG plays a role in the random migration of MEFs, we generated time series data for cells in which this protein was knocked down. The experimentally determined distributions for the Δx_‖_ and θ differed significantly from wildtype cells (Figs. 2C and D). Our computational analysis on the resulting cell tracks produced several insights into the role of RhoG in cell migration. Confidence-interval profiling of the predicted parameter values suggested that the predicted state 2 for WT and the RhoG KD are similar (i.e., the range of estimated parameter values for these states overlap) (**Supp. Fig. 5**). However, State 1 in the control case, in which the cells display strong directed migration, is replaced in the RhoG knock down case by a state in which the cells show little movement. Interestingly the time spent in these states is approximately the same in both cases. This leads us to hypothesize that RhoG plays a role in establishing persistent migration. A putative pathway for this mechanism is RhoG activation of Rac1 (via the DOCK180/ELMO) complex [19], [20]. However, whether this is the key pathway in this process, and how it is organized spatio-temporally, is a direction of future research.

### Inferring states from time series data

We next sought to determine if the two states corresponded to distinct phenotypes within the cell culture or if individual cells could transition between states. To test the possibility that individual cells change their migration state, we devised a method to predict the current state from individual cell tracks. Our approach uses a Bayesian prediction method based on the probability for a sequence of k successive steps arising from state 1 or state 2 (Methods).

Because RhoG is upstream of Rac1, we decided to make use of our existing FRET-based biosensor for Rac1 [21] to determine if the two predicted states correlate with differences in Rac1 activity. Ideally, we would have done this analysis in MEF cells. However, the high motility of MEFs makes it challenging to automatically track them for longer time periods within the field of view of the high magnification and high n.a. objective required for FRET biosensor imaging. Therefore, we decided to use slower moving HeLa cells. We ran our cell track analysis on HeLa cells containing the Rac1 biosenor (Fig. 3). Our method again predicted the presence of two states (Fig. 3A **insets**).

**Figure 3.**
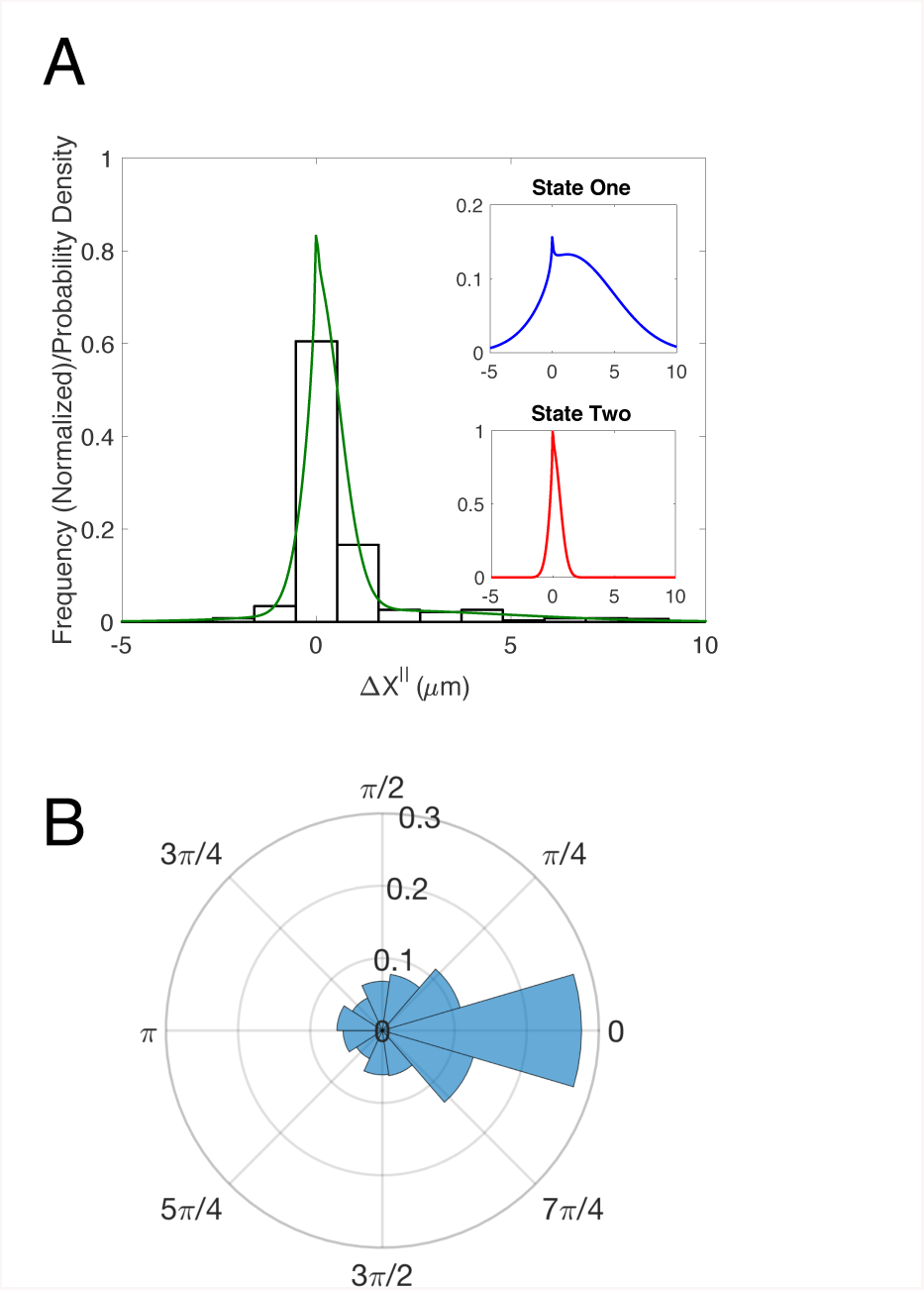
Fitting the model to HeLa cells expressing a Rac1 biosensor identified two states. **A** HeLa cells with Rac1 biosensor. Observed data (open bars), model fit (green) and individual pdfs for state one and state two (insets). **B** HeLa cells with Rac1 biosensor. Observed angular distribution for angle turned at each time step, *Δθ*.

Before running our state prediction methods on the experimental data, we first validated the approach using synthetic data. To generate this data, we performed computational simulations of the stochastic model using the parameter estimated from the experimental data. To generate cells tracks consistent with the experimental data, steps were generated in state 1 and 2 in proportion to *α* = 0.19 and 1 − *α*, respectively. For HeLa cells expressing the Rac1 biosensor this approach could correctly identify the states more than 90% of the time (**Suppl. Fig 7.**).

We next applied our method to the experimental data (Fig. 4). Our analysis suggested that individual HeLa cells do change their state of migration, with cells switching multiple times between states (Fig. 4A). To assess the biological relevance of the two states, we identified (see Supplementary Material for details) and counted the number of foci of Rac1 activity in each image and grouped these counts by the predicted state (Fig. 4B). Cells predicted be in state 1, which corresponds to the fast persistent state, had fewer Rac1 foci than those predicted to be in state 2 (Fig. 4B). The observed correlation between the predicted migration states and differences in Rac1 activity, provides further evidence to support the existence of distinct states of migration and the ability of cells to transition between these states.

**Figure 4.**
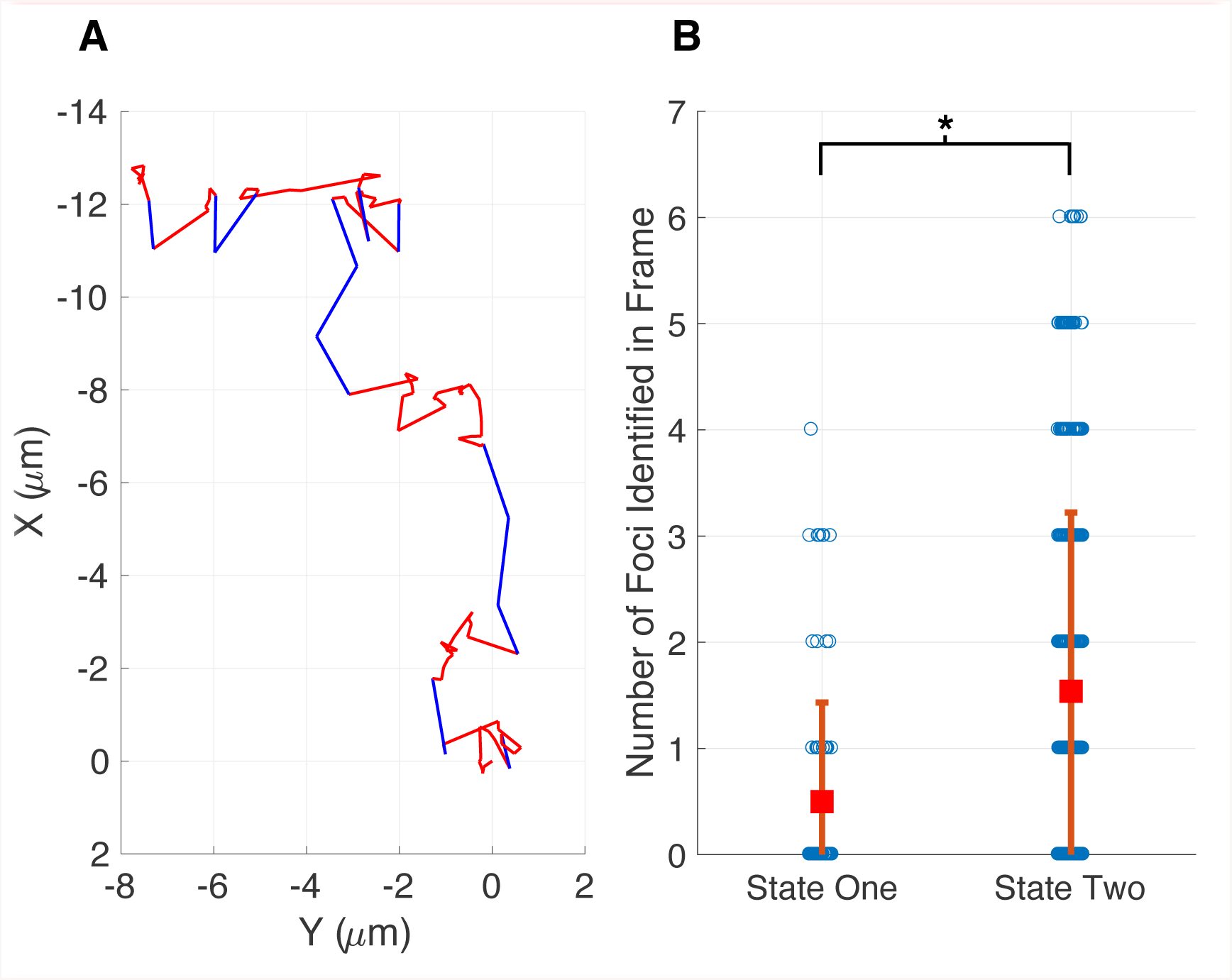
HeLa Cells expressing a Rac1 biosensor switch between states, and the states correspond with differential biological activity. **A** Example cell track showing switching between a fast persistent state (state one, blue) and a relatively stationary, turning, state (state two, red). States predicted from applying Bayes’ theorem with our model (Fig 3). **B** When in state two cells have a higher number of foci than when in state one. The probability for observing this effect by chance (if there was no difference between the states) is vanishingly small (supplementary figure 8).

## Discussion

We developed novel computational methods for analyzing the movement of randomly migrating cells. Our approach combines stochastic modeling with statistical inference methods to detect and quantify migratory phenotypes. Migrating cells have a biochemical, morphological, and structural orientation that persists as these cells move. Our model captures this ‘memory’ by conditioning the cell’s movement during the current time interval on its previous direction of motion. An important feature of our model is that analytic expressions for the probability densities for cell displacements parallel and perpendicular to the previous direction of motion can be found. This feature allows model parameters corresponding to cell velocity and persistence to be efficiently and accurately estimated from cell track data. We have validated all our approaches using simulated data, and then applied the methodology to study randomly migrating MEF and HeLa cells.

An important feature of our modeling approach is that it is general enough to allow for multiple states of migration. This feature allowed us to demonstrate that migrating cells randomly transition between modes of movement. Crucial to the detection of these states is the quantification of parameter values and the associated confidence in those estimates. This process allowed us to be confident in the existence of two states of migration for MEF cells and HeLa cells expressing a Rac1 biosensor.

The identification of multiple states of migration for MEF cells led us to assess the role of RhoG in establishing these states. To do this we used siRNA to reduce RhoG expression. This perturbation indicated that RhoG plays a role in orchestrating periods of persistent directed migration, a state that is lacking in cells where RhoG has been knocked down. To investigate the two states of migration in more detail, we developed a Bayesian approach to predict the current migration state of a cell from time series of the cell’s position. Using this method, we demonstrated that individual HeLa cells expressing a Rac1 biosensor switched between migratory states. Importantly, we were able to correlate these two states with differences in the distribution of Rac1 activity.

We believe that our methods provide useful tools for quantifying and characterizing cell migration. Our stochastic model characterizes cell migration using parameters with straightforward biological interpretations. Hence, application of this model can lead to biological insights not apparent in the data from visual inspection or simple quantitative measures. In this case, our analysis suggests a role of RhoG in orchestrating a persistent state of motion.

## Methods

### Computational methods

<u>Coordinate transformation.</u> We modeled cell migration as a stochastic sequence of steps characterized by the step size *r_i_* and directional angle *θ*_*i*_ (Fig. 1). Since we assume *r_i_* and *θ*_*i*_ to be realizations of independent random variables *R* and *Θ* the probability the cell moves (*r*, *θ*) is defined by

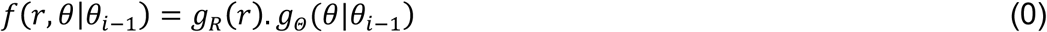

where *g_R_*(*r*) is the probability density function (pdf) for the step magnitude, which we take to have the normal distribution 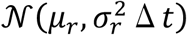 and 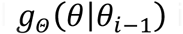 is the pdf generating the new orientation conditioned on the previous angle, which we take to have the normal distribution 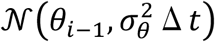
. The experimental data is collected in Cartesian coordinates (*X*, *Y*). In principle we could transform the data into the coordinates *R* and *Θ*. However this transformation cannot be completed uniquely, because there is no way to distinguish a backward step in which the cell maintains its direction of polarity 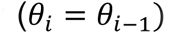 from one in which the front and back of the cell have reversed 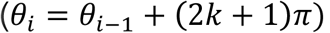. Furthermore the value of *θ*_*i*_ cannot be determined if *r*_*i*_ = 0. For these reasons, we transform the model to the coordinates 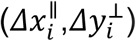, where these new variables correspond to changes in the centroid’s position during the i^th^ time interval that are parallel and perpendicular to the direction of the previous step.

To compare with the model the data needs to be manipulated to generate histograms for steps in the *x*^‖^ and *y*^⊥^ directions For each sequential triplet of coordinates 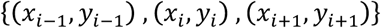, we rotate the steps as a rigid body about (*x*_*i*-1_, *y*_*i*-1_) by a four quadrant inverse tangent based on 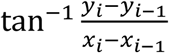. The result of this is that all steps are pre-orientated in a positive x-direction and initiated at (0,0) and can be plotted as histograms of step distance in the *x* and *y* direction: 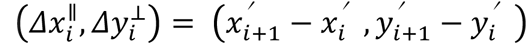.

The pdf for 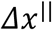 is: 
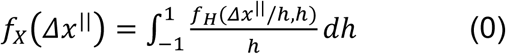
where *h* = cos (*θ*), and, 
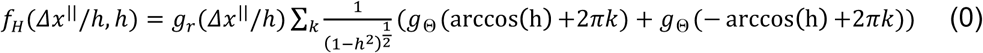

The expression for *f_Y_*(Δ*y*^⊥^) is similar, however now with 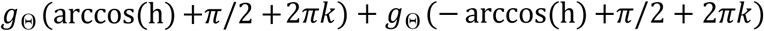 in the summation term. A derivation of these results is presented in the Supplemental Material.

<u>Parameter estimation.</u> Parameters were estimated by simulated annealing, which is a Monte Carlo method based on the Metropolis algorithm (24, 25). Initial choices of parameters generate an analytical solution (**Eq. 2**), which is scored against the experimental data 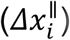 by the sum of least squared differences. At each step of the algorithm the parameters are updated by a small addition of Gaussian noise, if this update scores better than the current score then these parameters are accepted. If the score is higher, the parameter set is accepted with probability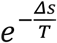, where *Δs* is the difference between the current and previous scores and *T* is the current temperature. Over the course of the fitting *T*, the temperature is reduced. This fixes the parameter choices into a local minimum. Here we choose a geometric cooling regime. Due to the stochastic nature of the simulation, and that there could be many local minima, it is necessary to run this fitting procedure multiple times. The best fit of this routine was then further refined using MATLABs fmincon routine, which was also used to assess the sensitivity of our fit to altering parameter values via confidence-interval profiling (**Sup. Fig 4**, Supplementary Materials for details).

The histograms were amalgamated from multiple cell tracks. For the case of two states, the pdf for *Δx* becomes

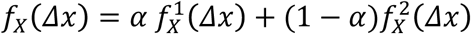

where *α* is the fraction of time spent in state 1 and the distributions 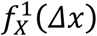 and 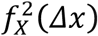 are parameterized by 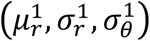 and 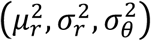, respectively.

Parameter sets were identified by multiple simulated annealing runs, followed by local-optimization routines.

<u>Validation of methods.</u> To validate the pdfs and the parameter estimation algorithm, we simulated cell tracks using the stochastic model (Fig. 1). Cell tracks were generated using two states, each with distinct parameter sets. At each step a state was chosen at random with probability 0.5. As above, the simulated cell tracks were used to construct the distributions fo 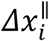 and 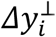. We assumed model parameters were not known and used the Monte Carlo method to fit **Eq. 2**., modified to two states (see below) to the simulated data for 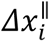. The Monte Carlo method quickly converged on the correct parameter values (**Suppl. Fig. 4**), validating the analytical solution to the model and our fitting procedure. In theory we also could fit the pdf for 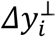. However, the pdf for 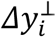 is symmetric, because there is no preferred direction of migration and therefore less informative than the distribution for 
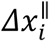. We found that we could maintain the accuracy of our parameter estimation while improving the computational cost by only considering the *Δx_i_*^‖^distribution. As a consistency check, we always verify that the estimated parameters accurately reproduce the pdfs for 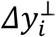 (**Suppl. Fig. 6**)

<u>State prediction.</u> To identify which state a cell is in at a given time, we used Bayes’ theorem to invert the problem. That is, we calculate the probability that a cell is in state *s_i_* given the experimental data. Note that in calculating this probability, we also get the false positive rate or p-value. To make a reliable prediction of *s_i_* may require an n-step window,where n is odd, such that, 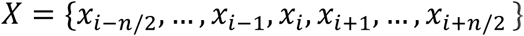. Then: 
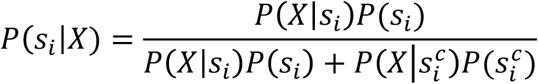
where *P*(*X*|*s_i_*) is calculated from the model, and we take *P*(*s_i_*) = *α*. Windows of length one, three and five were tested. For the case presented here, we found that the window of length one produced results similar to the other two window lengths.

<u>Foci identification.</u> Ratiometric images of the FRET based Rac1 biosensor were analyzed for localized regions of higher Rac1 activity near the periphery of the cell. We call these regions ‘foci’. We used custom application of the image processing toolbox in MATLAB to identify foci, which we define as contiguous regions within the cell that were simultaneously: (1) 60% above the average intensity of the cell, (2) greater than 100 pixels in area and (3) contained at least one point within 5 pixels of the cell edge. The length of time (or number of frames) that a cell could be followed for varied. So to not overweight any one cell, the number of image frames analyzed, *n*, was selected to maximize *n*×*m* where *m* is the number of cells with at least *n* images.

### Experimental methods

Cell tracks were generated through 20x phase contrast imaging of cells plated on fibronectin-coated coverslips. Cells were imaged using Ham’s F12K medium supplemented with 2% FBS for 18-24 hours at 5 minute intervals in a closed, heated chamber. 41 cells were tracked for at least 190 frames, obtained at 5 minute intervals.

IA32MEFs were transfected with either RhoG siRNA (CAGGTTTACCTAAGAGGCCAA) or Allstars Negative Control siRNA (Qiagen, USA). Control siRNA cells were incubated with 5uM CFDA green for 20min in serum-free DMEM. CFDA-labeled control cells were mixed with unlabeled RhoG siRNA cells immediately prior to the experiment. Cell tracks were generated through 10x DIC imaging of cells plated on 10ug/mL fibronectin-coated coverslips, using Ham’s F12K medium supplemented with 5% FBS. Images were acquired for at least 70 frames at 10 minute intervals, in a closed, heated chamber. This length of track was objectively identified as optimal by maximizing the total number of analyzed frames in the entire data set.

## Acknowledgements

This work was supported by Army Research Office grant W911NF-15-1-0631 to TE and KH, and NIH grants U01-EB018816 to TE and P41-EB002025 to KH.

## References

[1] Petrie R. J., Doyle A. D., and Yamada K. M., “Random versus directionally persistent cell migration,” Nature Publishing Group, vol. 10, no. 8, pp. 538–549, Jul. 2009.

[2] Cain R. J. and Ridley A. J., “Phosphoinositide 3-kinases in cell migration,” vol. 101, no. 1, pp. 13–29, Jan. 2012.

[3] Franca-Koh J., Kamimura Y., and Devreotes P. N., “Leading-edge research: PtdIns(3,4,5)P3 and directed migration,” Nat Cell Biol, vol. 9, no. 1, pp. 15–17, 2007.

[4] Finney A. C., Stokes K. Y., Pattillo C. B., and Orr A. W., “Integrin signaling in atherosclerosis,” Cell Mol Life Sci, vol. 74, no. 12, pp. 2263–2282, Feb. 2017.

[5] Lemarié C. A., Tharaux P.-L., and Lehoux S., “Extracellular matrix alterations in hypertensive vascular remodeling,” Journal of Molecular and Cellular Cardiology, vol. 48, no. 3, pp. 433–439, Mar. 2010.

[6] Hall A., “The cytoskeleton and cancer,” Cancer metastasis reviews, vol. 28, no. 1, pp. 5–14, Jun. 2009.

[7] Goley E. D. and Welch M. D., “The ARP2/3 complex: an actin nucleator comes of age,” Nat Rev Mol Cell Biol, vol. 7, no. 10, pp. 713–726, Oct. 2006.

[8] Ridley A. J., “Rho GTPases and actin dynamics in membrane protrusions and vesicle trafficking,” Trends Cell Biol, vol. 16, no. 10, pp. 522–529, 2006.

[9] Iden S. and Collard J. G., “Crosstalk between small GTPases and polarity proteins in cell polarization,” Nat Rev Mol Cell Biol, vol. 9, no. 11, pp. 846–859, Nov. 2008.

[10] Rottner K., Hall A., and Small J. V., “Interplay between rac and rho in the control of substrate dynamics,” Current Biology, vol. 9, no. 640, 1999.

[11] Ladwein M. and Rottner K., “On the Rho’d: the regulation of membrane protrusions by Rho-GTPases,” FEBS Lett, vol. 582, no.14, pp. 2066–2074, 2008.

[12] Jaffe A. B. and Hall A., “Rho GTPases: Biochemistry and Biology,” http://dx.doi.org.proxy1.athensams.net/10.1146/annurev.cellbio.21.020604.150721, vol. 21, no. 1, pp. 247–269, Oct. 2005.

[13] Dieterich P., Klages R., Preuss R., and Schwab A., “Anomalous dynamics of cell migration,” Proc Natl Acad Sci USA, vol. 105, no. 2, pp. 459–463, Jan. 2008.

[14] Othmer H. G., Dunbar S. R., and Alt W., “Models of dispersal in biological systems.,” Journal of mathematical biology, vol. 26, no. 3, pp. 263–298, 1988.

[15] Dimilla P. A., Quinn J. A., Albelda S. M., and Lauffenburger D. A., “Measurement of Individual Cell Migration Parameters for Human Tissue Cells,” AIChE Journal, vol. 38, no. 7, 1992.

[16] Rosello C., Ballet P., Planus E., and Tracqui P., “Model driven quantification of individual and collective cell migration,” Acta Biotheor, vol. 52, no. 4, pp. 343–363, 2004.

[17] Arrieumerlou C. and Meyer T., “A Local Coupling Model and Compass Parameter for Eukaryotic Chemotaxis,” Dev Cell, vol. 8, no. 2, pp. 215–227, Feb. 2005.

[18] Raue A., Kreutz C., Maiwald T., Bachmann J., Schilling M., Klingmüller U., and Timmer J., “Structural and practical identifiability analysis of partially observed dynamical models by exploiting the profile likelihood,” Bioinformatics, vol. 25, no. 15, pp. 1923–1929, Aug. 2009.

[19] Katoh H. and Negishi M., “RhoG activates Rac1 by direct interaction with the Dock180-binding protein Elmo.,” Nature, vol. 424, no. 6947, pp. 461–464, 2003.

[20] Katoh H., Hiramoto K., and Negishi M., “Activation of Rac1 by RhoG regulates cell migration.,” J Cell Sci, vol. 119, no. 1, pp. 56–65, Jan. 2006.

[21] Machacek M., Hodgson L., Welch C., Elliott H., Pertz O., Nalbant P., Abell A., Johnson G. L., Hahn K. M., and Danuser G., “Coordination of Rho GTPase activities during cell protrusion,” Nature, vol. 461, no. 7260, pp. 99–103, Mar. 2009.

